# The promotion of stress tolerant Symbiodiniaceae dominance in juveniles of two coral species due to simulated future conditions of ocean warming and acidification

**DOI:** 10.1101/2021.11.03.467090

**Authors:** Alyx P. Terrell, Emma Marangon, Nicole S. Webster, Ira Cooke, Kate M. Quigley

**Affiliations:** Australian Institute of Marine Science, Townsville, QLD, Australia; College of Science and Engineering, James Cook University, Townsville, QLD, Australia; AIMS@JCU, Townsville, QLD, Australia; Australian Centre for Ecogenomics, University of Queensland, Brisbane, Australia; Australian Antarctic Division, Kingston, Tasmania, Australia; Department of Molecular and Cell Biology, James Cook University, Townsville, QLD 4811, Australia; Centre for Tropical Bioinformatics and Molecular Biology, James Cook University, Townsville, QLD 4811, Australia

## Abstract

The symbiotic relationship between coral and its endosymbiotic algae, Symbiodiniaceae, greatly influences the hosts’ potential to withstand environmental stress. To date, the effects of climate change on this relationship has primarily focused on adult corals. Uncovering the effects of environmental stress on the establishment and development of this symbiosis in early life stages is critical for predicting how corals may respond to climate change. To determine the impacts of future climate projections on the establishment of symbionts in juvenile corals, ITS2 amplicon sequencing of single coral juveniles was applied to *Goniastrea retiformis* and *Acropora millepora* before and after exposure to three climate conditions of varying temperature and *p*CO_2_ levels (current and RCP8.5 in 2050 and 2100). Compared to ambient conditions, juvenile corals experienced shuffling in the relative abundance of *Cladocopium* (C1m, reduction) to *Durusdinium* (D1 and D1a, increase) over time. We calculated a novel risk metric incorporating functional redundancy and likelihood of impact on host physiology to identify the loss of D1a as a ‘low risk’ to the coral compared to the loss of “higher risk” taxa like D1 and C1m. Although the increase in stress tolerant *Durusdinium* under future warming was encouraging for *A. millepora*, by 2100, *G. retiformis* communities displayed signs of symbiosis de-regulation, suggesting this acclimatory mechanism may have species-specific thresholds. These results emphasize the need for understanding of long-term effects of climate change induced stress on coral juveniles and their potential for increased acclimation to heat tolerance through changes in symbiosis.

**Originality Statement:** Here we assessed changes in the uptake and establishment of Symbiodiniaceae in the early lifehistory stages of two coral species under future climate scenarios. Our study represents the first such assessment of future climate change projections (increased temperature and *p*CO_2_) influencing Symbiodiniaceae acquisition and specifically shows a community structure dominated by the stress tolerant genus *Durusdinium*. We also develop a novel risk metric that includes taxonomic function and redundancy to estimate the impact of symbiont taxa changes on coral physiology. Through the risk metric, we relate the stress-induced changes in symbiont community structure to the likelihood of functional loss to better understand the extent to which these changes may lead to a decrease in coral health.

## Introduction

Increasing global temperatures due to anthropogenic climate change is the greatest current threat to corals, resulting in three global mass bleaching events since the 1980s (1998, 2010, 2014-2017), with bleaching becoming more prevalent and extreme (Hughes *et al*., 2017). Coral bleaching occurs when endosymbiotic dinoflagellates (family Symbiodiniaceae) are lost from the host coral tissue. As Symbiodiniaceae support their host’s energy demands through the provision of photosynthates, the breakdown of this relationship may lead to coral mortality (Brown, 1997, Suggett *et al*., 2017). Due to high rates of ocean warming and increased partial pressure of carbon dioxide (*p*CO_2_), there is grave concern that the incidence of bleaching events may be too rapid to allow for natural adaptation to occur (Hoegh-Guldberg *et al*., 2007, Hughes *et al*., 2017). Understanding the effects of climate change on coral-Symbiodiniaceae interactions is therefore critical for predicting coral reef resilience to future climate.

Due to the critical role of the endosymbiotic Symbiodiniaceae in energy translocation within the coral host, there has been an emphasis on increased taxonomic resolution of the Symbiodiniaceae present in the host using genetic markers such as the internal transcribed spacer 2 (ITS2) region (Pochon *et al*., 2014). Symbiodiniaceae are comprised of 7 currently identified genera (formerly 9 clades), each of which is subset into species (formerly ‘types’) (LaJeunesse *et al*., 2018). Host environmental sensitivity differs according to the predominant algal symbiont or community. In adults of *Acropora millepora*, for example, changes in Symbiodiniaceae relative abundance towards *Durusdinium* (formerly clade D) may increase host heat tolerance by 1–1.5 °C (Berkelmans & van Oppen, 2006). Therefore, the dominance of specific Symbiodiniaceae genera within the coral host can promote greater resilience under environmental stressors such as increased sea surface temperature or to other stressors present in the environment (Baker *et al*., 2004, Berkelmans & van Oppen, 2006, Oliver & Palumbi, 2011, Stat & Gates, 2011, Stat *et al*., 2013).

While shuffling of specific symbiont taxa may improve adult coral tolerance to stress, it is less understood if symbiont shuffling occurs in coral early life-history stages. Coral juveniles generally establish their initial endosymbiont community either through direct transmission from the maternal colony (vertical transmission), or via environmental acquisition (horizontal transmission) (Baird *et al*., 2009) or a combination of the two (mixed-mode transmission) (Quigley *et al*., 2018). Vertical symbiont establishment provides the opportunity for larvae and subsequent juveniles to acquire a potentially advantageous community of endosymbionts best suited for the local, maternal environment. Conversely, horizontal transmission allows for more flexibility to acquire novel symbionts that could be advantageous under changing environmental conditions or after large dispersal distances. For example, selecting for *Durusdinium* species promoted greater survival rates in juvenile *A. tenuis* when exposed to increased temperature and light levels (Yuyama *et al*., 2016).

Different Symbiodiniaceae taxa also provide different host benefits. For example, *Durusdinium* may provide increased survival to juveniles under simulated bleaching events (Quigley *et al*., 2020). Further, growth rates vary in juveniles between those dominated by either *Cladocopium* or *Durusdinium* (Cantin *et al*., 2009, Yuyama & Higuchi, 2014, Quigley *et al*., 2020), linked to differences in carbon translocation between genera (Cantin *et al*., 2009). Physiological differences also exist at finer taxonomic scales, for example, within *Durusdinium* D1a compared to D1, driven by differences in physiology, like variable photosystem II activity (Suggett *et al*., 2015).

The diversity of the symbiont community during coral juvenile development may also be important for growth and survival. There is some evidence that the acquisition of diverse endosymbionts may promote higher growth rates (Yuyama & Higuchi, 2014) and increase survival (Suggett *et al*., 2017). Compared to adult corals, early life-history stages generally have more diverse endosymbiont communities (Quigley *et al*., 2017), which eventually winnow to a more stable, lower diversity community (Lee *et al*., 2016, Rouzé *et al*., 2019). In some cases, symbiont communities in juvenile corals may be impacted by environmental factors such as increased temperature and light (Abrego *et al*., 2012), *p*CO_2_ (Suwa *et al*., 2010) or completely halted by increased temperatures with minimal impacts of *p*CO_2_ (Sun *et al*., 2020). Moreover, the acquisition of symbionts during coral early life-history stages has been assessed only for a limited number of coral species (Cantin *et al*., 2009, Yorifuji *et al*., 2017, Quigley *et al*., 2019, Quigley *et al*., 2020), and the effects of elevated temperature and *p*CO_2_ on this process are poorly understood. Combined, these results highlight the importance of Symbiodiniaceae community diversity for the fitness of coral adult and early life-history stages and highlight an increased need for a better understanding of fine-scale host-symbiont interactions.

To assess changes in Symbiodiniaceae acquisition during coral developmental stages under future climate scenarios, two coral species, *Goniastrea retiformis* and *Acropora millepora*, were exposed to varying levels of temperature and *p*CO_2_ predicted for the years 2050 and 2100 under RCP 8.5 and compared to present day levels. Juveniles were sampled and the ITS2 region of the symbionts was targeted using amplicon sequencing. The dynamics of symbiont community changes (prevalence, relative abundance and diversity) were assessed under exposure to these climate treatments over time (0, 10 days and 4-5 weeks) in coral juveniles from these two species.

## Results and Discussion

### Coral juveniles reared under future climate scenarios

A full description of the experimental design, DNA extractions and sequencing pipeline can be found in the Supplementary methods. Briefly, in October 2017, gravid colonies of *G. retiformis* and *A. millepora* were collected from Geoffrey Bay (Great Barrier Reef, Australia) and transported to holding mesocosm tanks at the National Sea Simulator (SeaSim) at the Australian Institute of Marine Science as per details outlined in (Uthicke *et al*., 2020). Corals were acclimated to the following conditions (~27°C, *p*CO_2_ 400±60ppm) until spawning (*G. retiformis*: 8^th^ of November 2017; *A. millepora*: 12^th^ December 2017). After spawning, larvae were reared in mass culture tanks for 3-4 weeks. Larvae were then distributed to 6-well plates filled with filtered seawater containing autoclaved crustose coralline algae (CCA) to induce larval settlement. At the first sampling time point of newly settled recruits (T0), individuals were sampled and preserved in absolute ethanol in Eppendorf tubes (n = 3 individuals per tube). Following initial sampling, 2 × 6-well plates were placed in outdoor mesocosm tanks (n = 3 tank replicates per treatment) representing ambient (present day ~28.5°C, *p*CO_2_ 400±60ppm), 2050 (+1°C offset, *p*CO_2_ 685±60ppm), and 2100 (+2°C offset,*p*CO_2_ 940±60ppm) conditions forecast under 8.5 RCP (Meinshausen *et al*., 2011, Collins *et al*., 2013) (Supplementary Table 1). Sampling was repeated across each tank after 10 days (T1) and 4 (*G. retiformis*) or 5 (*A. millepora*) weeks (T2) of exposure to climate conditions. All samples were stored at −80°C prior to DNA extraction and ITS2 amplicon sequencing of Symbiodiniaceae communities (see Supplementary material for information about the challenges of Symbiodiniaceae taxonomy and the use of taxonomic names compared to ASV designations). However, due to lack of gel electrophoresis bands present in T0 and T1 *A. millepora* samples following PCR amplification, these time points were not sequenced.

### Changes in relative abundance of *Goniastrea retiformis* juveniles

The ambient treatment in this experiment represented average reef conditions over the past ~20 years, allowing for comparisons of symbiont acquisition under elevated temperatures and *p*CO_2_ expected by 2050 and 2100 under RCP8.5. Corals generally live close to their thermal maximums, and +1 and +2 degrees of temperature increases impact coral offspring physiology and that of their symbionts (Abrego *et al*., 2012), suggesting that these treatments represent a “stress” for coral offspring.

Adult *G. retiformis* are typically dominated by Symbiodiniaceae of the genus *Cladocopium* (Leveque *et al*., 2019), however studies conducted on other species propose that under elevated temperature, an increase in more thermally tolerant *Durusdinium* would be expected (Baker *et al*., 2004, Berkelmans & van Oppen, 2006, Stat *et al*., 2013). Similar to adults, *G. retiformis* juveniles in this study were typically dominated by *Cladocopium*, but only at T0, although *Cladocopium* abundances remained higher in juveniles through time under ambient conditions compared to the 2050 and 2100 treatments (Fig. 1). At T1 and T2, *Durusdinium* were found at higher relative abundance, especially in the 2050 treatment, accompanied by a decrease in *Cladocopium* relative abundance. This pattern was evident in both the T1 and T2 timepoints, which may be due to the extended duration of stress or juvenile ontogeny and winnowing of symbiont communities. Interestingly, as the average relative abundance of *Cladocopium* decreased through time, this genus was replaced by *Durusdinium* or other taxa (e.g., B1g, I4) in 2050 and 2100 scenarios at T2 (Fig. 1). This could suggest that *G. retiformis* has the ability to shuffle their symbiont communities towards more heat-resistant taxa in the “shorter” term (up to 2050), taking on the classic shuffling pattern between *Cladocopium* to *Durusdinium*, but that future temperature and *p*CO_2_ conditions projected for 2100 may represent a threshold in which this acclimatory shuffling mechanism breaks-down, manifesting as the lowered relative abundances of *Cladocopium* and *Durusdinium* and increases in potentially “non-symbiotic” taxa.

**Fig. 1.**
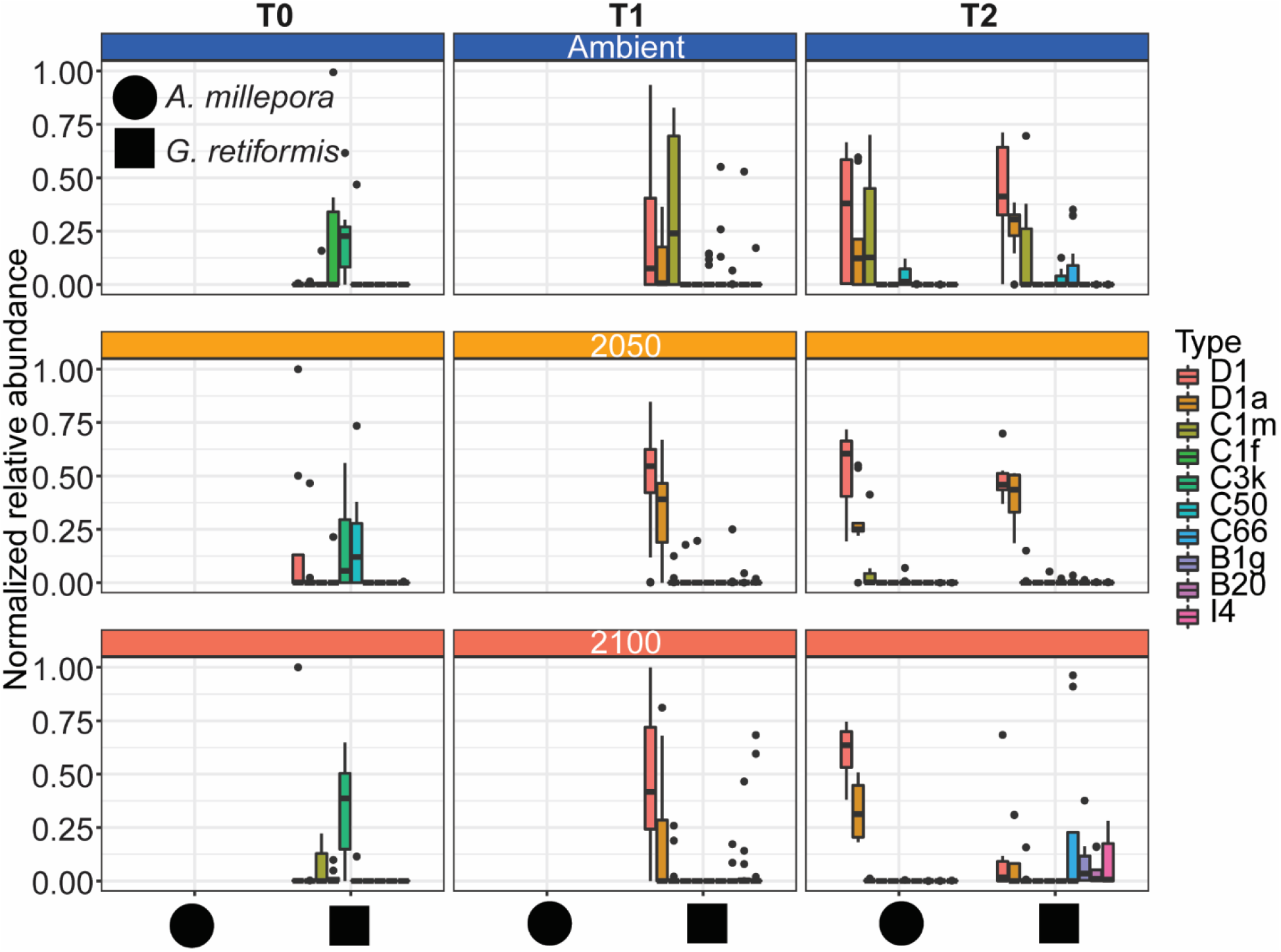
Effects of elevated temperature and *p*CO_2_ on Symbiodiniaceae communities in coral juveniles of *Acropora millepora* and *Goniastrea retiformis*. Gravid colonies of *A. millepora* and *G. retiformis* collected in the wild were spawned, gametes were produced and settled into recruits in outdoor tanks across three experimental treatments with three replicate tanks each: present-day (ambient, 28.5°C, *p*CO_2_ 400±60ppm), 2050 (+1°C offset, *p*CO_2_ 685±60ppm), and 2100 (+2°C offset, *p*CO_2_ 940±60ppm). Samples were collected at 14 days post-settlement (T0), 10 days (T1) and four weeks of treatment exposure (T2). DNA was extracted from individual recruits. Following PCR amplification and amplicon sequencing of the ITS2 region targeting Symbiodiniaceae. For *A. millepora*, only samples collected at T2 were sequenced. Bioinformatic analyses were performed, and Amplicon Sequence Variants (ASVs) identified. ASVs were variance normalized and relative abundances calculated. Here show the ten most abundant Symbiodiniaceae taxa (at a >4.2% relative abundance cut-off) across each juvenile. Overall, we observed shifts in symbiont taxa in response to time (for *G. retiformis*) and by treatment. Outliers were not removed.

*G. retiformis* juveniles sampled from the ambient treatment at T0 (i.e. before recruits were exposed to treatment conditions) were dominated by *Cladocopium* (as discussed above), but more specifically, by C3k and C1f (relative abundance >10% in >42% of individuals) and other background taxa including C1m, C50, and C3.14 (relative abundance >10% in >1% per juvenile) (Fig. 1). Within this ambient treatment, there were changes in the relative abundance at this initial timepoint to increased *Durusdinium* at T1 and T2, which may represent first initial uptake or rearing conditions before treatments were applied. Symbiodiniaceae shifts were observed in the T1 ambient treatment, specifically with the dominance of C1m (relative abundance >10% in 55% of individuals), D1 (relative abundance >10% in 50% of individuals), D1a (relative abundance >10% in 33% of individuals), and C66 and C50 (relative abundance >10% in 16% of individuals). By four weeks under ambient conditions, juveniles were dominated by D1 and D1a (relative abundance >10% in 95% of individuals), C1m (relative abundance >10% in 40% of individuals), and C66 (relative abundance >10% in 25% of individuals) (Fig. 1). Before exposure to 2050 and 2100 conditions (at T0), juveniles were dominated by C50 (relative abundance >10% in 62% of individuals), C3k (relative abundance >10% in 50% of individuals), D1 (relative abundance >10% in 25% of individuals), and very low relative abundances of D1a, C1p, C1f, and C29 (relative abundance >10% in 12% of individuals) (Fig. 1). After 10 days under 2050 and 2100 treatments at T1 and the 2050 treatment at T2, *Cladocopium* relative abundances dropped. By T2 in 2100, background types represented the highest relative abundance (relative abundance >10% in 63% of individuals), including D1.1, I4 and unidentified types (relative abundance >10% in 38% of individuals). The change toward *Durusdinium*-dominance may be driven by juvenile age (or duration of stress exposure), although climate treatment was the main driver of the change in individual Amplicon Sequence Variants (ASVs). A total of 70 ASVs (grouped by “species” designations in Fig. 2), significantly changed in abundance (generalized linear models, see Supplementary information), all of which changed in relation to treatment or treatment*species (Fig. 2). Changes by species and time were only related to changes in ASV abundance with the climate treatment interaction. This suggests that while juvenile age and length of climate exposure may result in changes in the relative abundance of ASVs with the Symbiodiniaceae communities, climate-related pressures were the main driver of those changes in juvenile corals during early development.

**Fig. 2.**
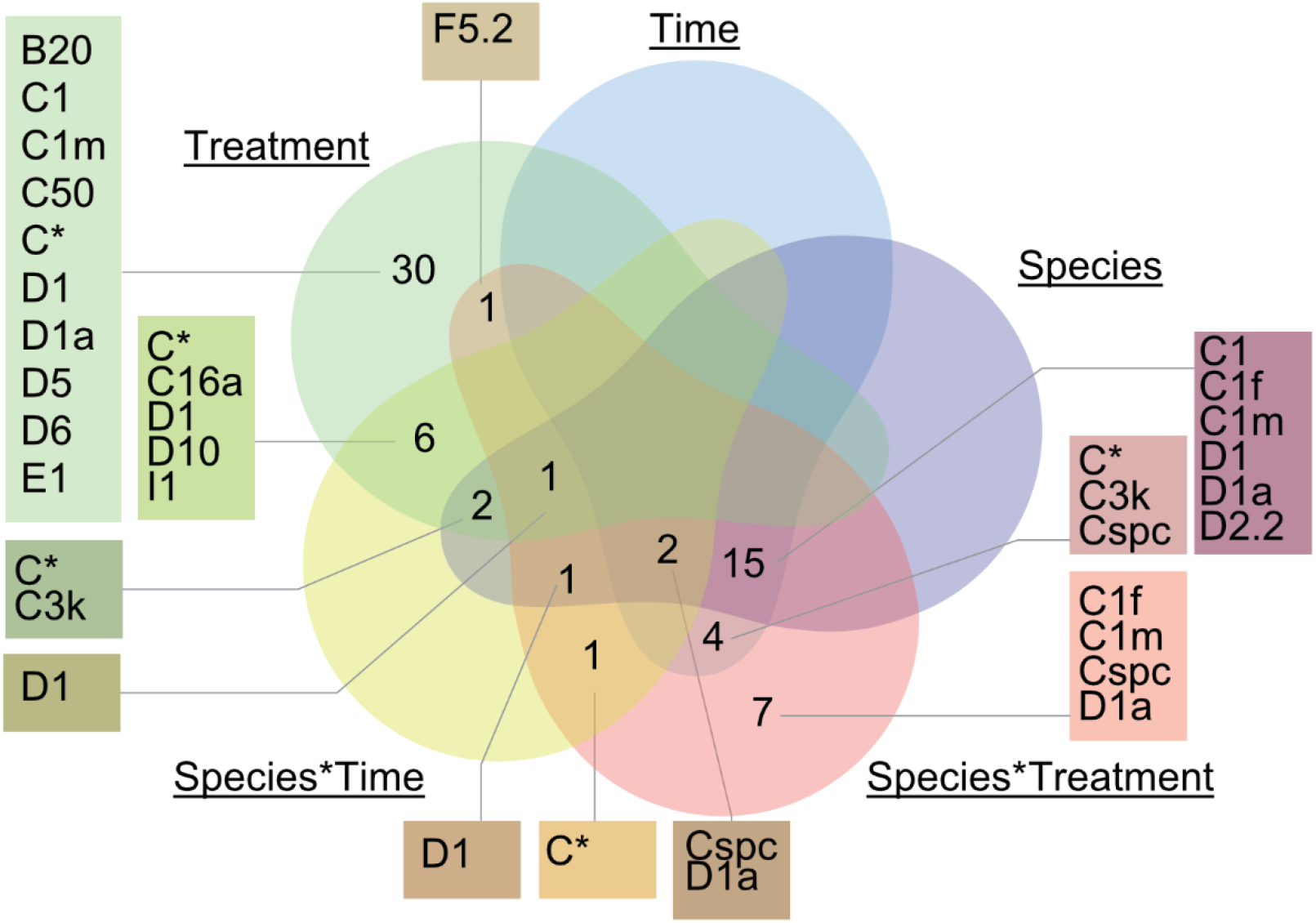
Venn diagram illustrating ASVs that significantly changed in relative abundance between Treatments/Time/Species and their interactions. A generalized linear model was run to determine which factors (time, treatment, species, or interactions between) were linked with the greatest number of ASVs that significantly changed in relative abundance using the package DESeq and DESeq2 (v. 1.26.0). It is worth noting that Species*Time corresponds to *A. millepora* T2 vs. *G. retiformis* T2 and Species to *A. millepora* (T2) vs *G. retiformis* (T0, T1, T2). Sequences were aligned to the EMBL-EBI web server program of Clustal Omega to identify the closest matched ITS2 taxa. There were no ASVs that were significantly associated with the Time*Treatment interaction and are not listed in the figure.

### Changes in relative abundance of *Acropora millepora* juveniles

*Acropora millepora* juveniles, which were sequenced only at five weeks (T2), were dominated by *Durusdinium* across treatments. *Cladocopium* was present but decreased in relative abundances with increasing climate stress (Fig. 1). This is consistent with other studies in *A. millepora* juveniles showing a change towards *Durusdinium*-dominance at higher temperatures (Abrego *et al*., 2012) as well as in adult corals of the same species that showed an increased abundance of *Durusdinium* when corals are exposed to elevated sea surface temperatures and/or human disturbances (Oliver & Palumbi, 2011, Chen *et al*., 2019, Claar *et al*., 2020). Although community changes towards *Durusdinium* may increase host thermal tolerance, it may concomitantly decrease growth rates in *Durusdinium*-dominated *A. millepora* juveniles (Little *et al*., 2004, Cantin *et al*., 2009). Therefore, there may be limits to the benefits of hosting *Durusdinium* for the coral *A. millepora*. Specifically, in this study, juveniles of *A. millepora* were dominated by symbionts from D1 and D1a across all climate treatments (relative abundance >10% in >80% of sample, Fig. 1), including 2050 and 2100 (>10% relative abundance) and in 66% of ambient juveniles (>10%). Other background types like C1m, C50 and D5 were also present at low relative abundances across treatments. C1m was abundant in ambient conditions (>10% relative abundance in 55% of samples) but decreased in relative abundance with increasing stress levels (2050: >10% relative abundance in 10% of samples, 2100: >10% relative abundance in 0% of samples). Likewise, other background types present below threshold values were more abundant in the ambient treatment juveniles compared to those of 2050 and 2100, with 2100 samples comprising very few background types even below the abundance threshold. As seen in *G. retiformis* juveniles, the increased relative abundance of D1 and D1a in conjunction with the decreased abundance of background types in both 2050 and 2100 treatments may suggest the ability of *A. millepora* juveniles to acclimate to future climate conditions for at least short experimental periods. This demonstrated potential, if applicable in the wild, may promote greater survival under future climate scenarios, thereby increasing the opportunity for coral survival through natural regenerative events, like coral spawning and subsequent recruitment.

### Shifts in the diversity of Symbiodiniaceae occurred across time, treatments, and coral species

Changes in the symbiont community in coral early life history stages have been observed in both field and lab-based experiments (Abrego *et al*., 2009, Quigley *et al*., 2017, Quigley *et al*., 2019), in which the overall diversity as well as individual symbiont taxa can strongly influence the host across traits such as growth, survival and thermotolerance (Mieog *et al*., 2009). Endosymbiotic community diversity may vary in response to environmental conditions, including temperature and nutrients (Gong *et al*., 2018). In our study, climate treatments did not impact diversity (Shannon index) within each species (*p* > 001; Fig. 3; Supplementary information). However, diversity in *G. retiformis* juveniles changed significantly over time, with an overall lower diversity at T1 compared to T2 (ANOVA: *F*_2,97_ = 11.2, *p* < 0.01). When comparing diversity between species at T2, mean diversity was higher in *A. millepora* compared to *G. retiformis* in juveniles under 2100 conditions (ANOVA, Species: *F*_1,54_ = 6.0, *p* = 0.02; Species*Treatment: *F*_2,54_ = 3.4, *p* = 0.04; *G. retiformis* 2100 - *A. millepora* 2100: *p* = 0.03).

**Fig. 3.**
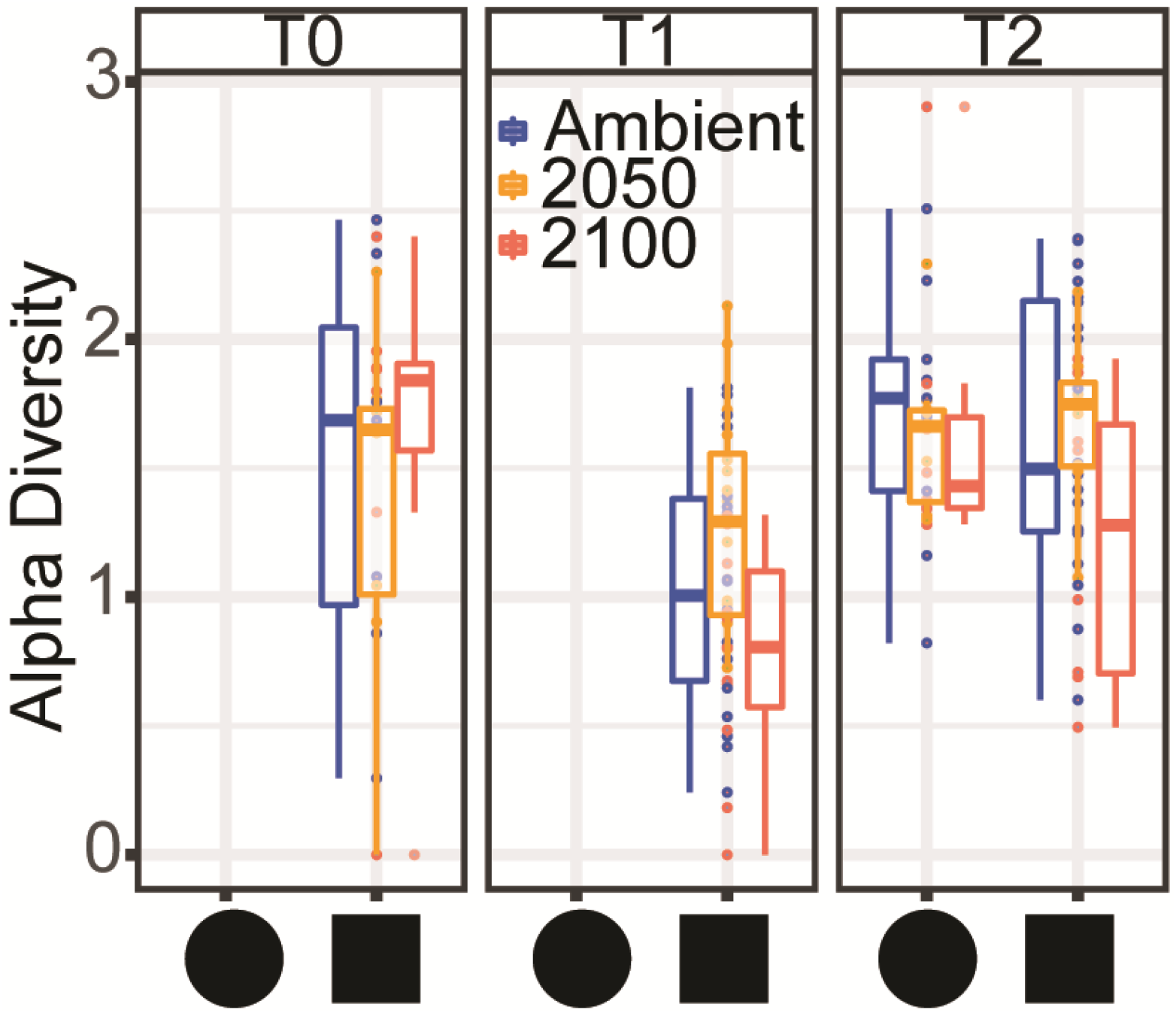
Shannon diversity index in *A. millepora* and *G. retiformis* juveniles across temperature and *p*CO_2_ treatments (ambient, 2050, 2100) and time (T0, T1, T2). Shannon diversity metric was calculated using normalized ASVs abundances calculated for each individual juvenile using the package Phyloseq (version 1.30.0) in R. *A. millepora* was only sequenced at timepoint T2.

Taken together, these results suggest that at least at the final timepoint, Symbiodiniaceae communities in *A. millepora* were more diverse and evenly distributed across individual juveniles compared to *G. retiformis* under high environmental stress, moving towards a community involving *Durusdinium*. Considering that an increased dominance of *Durusdinium* was observed under stress (Fig. 1), the unaffected diversity under high temperatures and *p*CO_2_ may suggest that this taxa has the ability to outcompete other symbiont taxa.

### Changes in specific Symbiodiniaceae taxa driven by species-specific responses to temperature and *p*CO_2_

*Durusdinium* are of particular ecological interest due to their relative tolerance to stress compared to other symbionts (Fabricius *et al*., 2004, Morikawa & Palumbi, 2019). In accordance with our data showing the increased abundances of this genus through time and in stress treatments, we found that the taxa most likely to change were those that were identified to *Durusdinium* ASVs, including *D. glynnii* (15 ASVs to D1) and *D. trenchii* (9 ASVs to D1a). This change may correspond to changes in the coral host from *Cladocopium*-dominance, leaving available niche space for *Durusdinium*-dominance through time. Shuffling was also pervasive in *Cladocopium*, including a total of 12 *Cladocopium* ASVs shuffling across our treatment conditions (Fig. 2). The most common taxa that shuffled included C50 (4 ASVs), C1 (3 ASVs), and C1m (3 ASVs), mostly driven by climate treatment effects (Fig. 2). Interestingly, *G. retiformis* juveniles sampled from the 2100 treatment group shuffled over time from C50 (eight ASVs in T0 and T1) to C66 dominance (one ASV in T2 detected through outlier analysis). However, changes in C50 were more directly related to treatment effects whilst C66 changed by treatment*species interaction (Fig. 2). While *Cladocopium* C66 has not been previously identified in *Goniastrea* adults (Leveque *et al*., 2019), it has been found at low abundance in corals (LaJeunesse *et al*., 2003). However, C66 has been reported from *A. tenuis* juveniles, with an increase of C66 over 71 days in the wild (Quigley *et al*., 2019). This suggests that our T2 treatment may represent the initial establishment of a *Cladocopium* species containing the C66 marker and not an experimental artifact.

### The loss of specific Symbiodiniaceae taxa impact coral host functions

Symbiodiniaceae taxa are functionally diverse (Suggett *et al*., 2015, Swain *et al*., 2017), with different taxa performing diverse functions within the coral host (Muscatine, 1990). The loss or gain of particular taxa could, as a result, cause physiological changes in host functioning, including in nutrient transfer (e.g. carbon translocation) or transfer of stress tolerance to the host. To infer potential consequences on host functioning due to the taxonomic changes measured here, we developed a risk metric that incorporated the top 16 of the most highly abundant symbiont taxa in our juveniles, their functional roles, and the risk accompanied by the loss of taxa relative to the redundancy of their functional roles (see Supplementary material for full methods, Fig. 4). Based on the metric R_k_ (*sensu* Swain *et al*., 2017), Supplementary Table 4), where a higher R_k_ value is indicative of increased thermal tolerance, *Cladocopium* taxa vary in their sensitivity to high temperatures. For example, *Cladocopium goreaui* (C1) are more thermally-sensitive (R_k_ = 21.72) compared to C40 (R_k_ = 55.32). When combined with relative abundance, taxa like C40 experienced dramatic changes in relative abundance (R_k_ shift = 55) between ambient and climate stress treatments. Redundancy within juveniles for this taxon was also relatively low (redundancy rank = 6), resulting in the assignment of this taxa to a “medium risk” category if lost (Fig. 4). This suggests that given C40 had relatively high heat tolerance, thereby providing the coral host with potentially greater stress resilience, the loss of this symbiont could be significant to host health given its low functional redundancy. *Durusdinium trenchii* (D1a), also generally known as a stress tolerant symbiont, had a correspondingly high R_k_ (37.03) and experienced relatively large changes in relative abundance between treatments. However, functional redundancy was high (~13), suggesting that there is a diminished risk to host health if this taxon is lost. Importantly, taxa like C1, C1b, and D4 have low thermal tolerance, experienced large changes in relative abundance, and had low functional redundancy, suggesting the loss of these taxa are of greater risk to the host. Combined, this novel risk framework suggests that the loss or gain of different symbionts may not be functionally equivalent across these simulated future climate treatments in which each loss of particular symbionts may impact the host through downstream impacts on host function. These outcomes may also be species-dependent, where some coral species may rely more heavily upon specific Symbiodiniaceae taxa to fulfill certain functions.

**Fig. 4.**
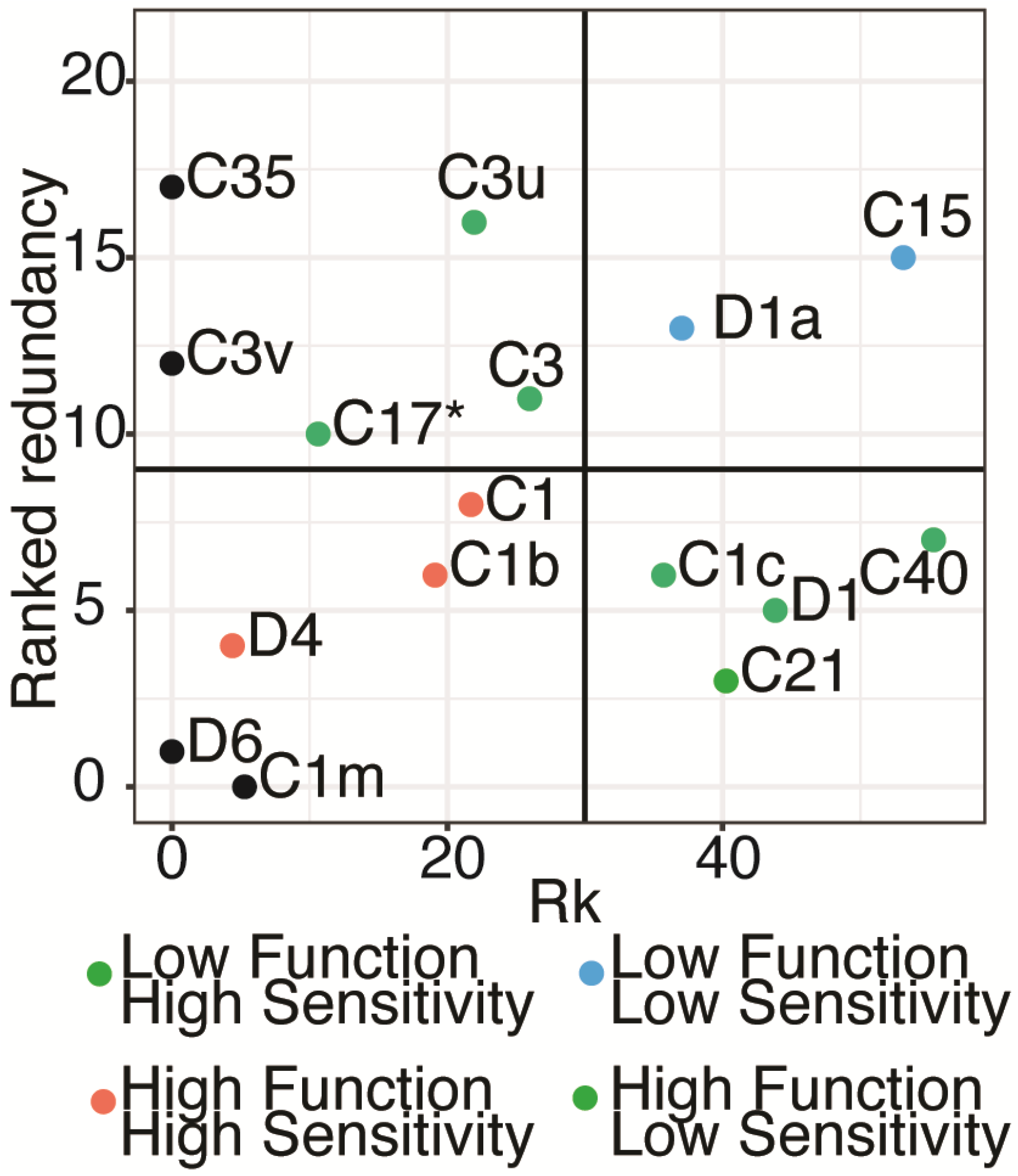
Risk metric describing the breakdown of host functioning due to Symbiodiniaceae loss. The metric included 16 of the most highly abundant Symbiodiniaceae ASVs and included rankings of the likelihood of shifts (horizontal axis; Rk score) and functional redundancy of each taxon. Rk values are calculated based on the IPRAPA-based Rk values in Swain et al. (2017) which measure the thermo-sensitivity of taxa. Redundancy rankings were calculated based on 12 health-associated measurements (e.g. Fv/Fm, chlorophyll content, lipid assessment) derived from 17 published experimental studies (see Supplementary). For each taxa, measurements were averaged across all studies per trait, with each function per taxa ranked from smallest to largest change, where a small change is indicative of greater redundancy and potential resilience of the host against stress. Final rankings were scored using a Borda Rank method and averaged to result in a relative ranking per function per taxa (from 1: important taxa for function – 21: less important taxa for function). Based on these two metrics, taxa were broken up into High (red points), Medium (green points), and Low (blue points) risk categories. For example, taxa in the High risk (lower left quadrat) identifies taxa which lost would have the greatest risk to impact host functioning due to their low redundancy and high functional importance. Taxa placed along the axis and colored in black were ranked based on only one value due to lack of data for the calculation of the Rk value.

## Conclusion

Overall, we observed an increase in relative abundance of *Durusdinium* over time in relation to juvenile age and duration of climate stress exposure. In addition, we found a significant capacity for coral juveniles to take up D1 and D1a symbionts at treatments with increasing temperature and *p*CO_2_, which generally corresponded with a decrease in *Cladocopium*. These results support the idea of ‘shuffling’ algal symbiont communities for increased acclimation to stress, as seen in studies of adult corals. Longer-term observations of coral juveniles are required to understand the ontogenic consequences of temperature and *p*CO_2_ on symbiosis as it may provide coral offspring with an increased capacity for heat tolerance and survival.

## Acknowledgments

The authors declare no conflict of interest. Sequencing data is available under BioProject PRJXXX in NCBI. This study was part of the Evolution21 project funded by AIMS and would like to thank the AIMS SeaSim staff and volunteers helping collect the colonies (permit G13/36318.1), spawn the corals, and carry-out the husbandry and experimental set-up and maintenance. The authors acknowledge the Gurambilbarra Wulgurukaba Traditional Owners of Magnetic Island, and pay our respects to their Elders, past, present, and emerging.

## Supporting Information

Supplementary material

