## Supplementary Methods for "The promotion of stress tolerant Symbiodiniaceae dominance in juveniles of two coral species due to simulated future conditions of ocean warming and acidification"

### A. Species selection

The common scleractinian corals *Goniastrea retiformis* and *Acropora millepora* were selected as model species for our experimental study. *G. retiformis* is a horizontally transmitting coral primarily acquiring its symbiont partners from the environment (Baird *et al.*, 2009) This coral species is associated with Symbiodiniaceae of the genus *Cladocopium* (formerly known as Clade C) as well as the heat-tolerant genus *Durusdinium* (formerly Clade D), which likely contributes to the host high thermal tolerance (Darling *et al.*, 2012, Leveque *et al.*, 2019). *Acropora millepora*, is another horizontally transmitting coral, primarily associated with *Durusdinium* but also harboring *Cladocopium* in low abundances (Quigley *et al.*, 2020). Despite the association with *Durusdinium*, *A. millepora* is a stress-sensitive species (Darling *et al.*, 2012).

### B. Coral collection, spawning, larval rearing and juvenile settlement

In October 2017, gravid colonies of *Goniastrea retiformis* and *Acropora millepora* were collected from Geoffrey Bay, Magnetic Island (S 19°09.326′, E 146°51.861′; Great Barrier Reef Marine Park Authority Permit Number: G13/36318.1). Colonies were transported to the National Sea Simulator at the Australian Institute of Marine Science (AIMS, Townsville, Queensland) where they were maintained in outdoor tanks until spawning (*G.* retiformis: 8$^{th}$ of November 2017; *A. millepora*: 12$^{th}$ December 2017). Following spawning, gametes were collected and mixed for fertilization. After rearing larval cultures in tanks for 3-4 weeks, larvae were randomly distributed across sterile 6-well plates (n=18 per species; n=20 larvae per well) filled with filtered seawater (0.04 µm) and containing autoclaved crustose coralline algae (CCA) to induce larval settlement. Plates were maintained at constant temperature (~27.5°C) and seawater was replaced daily.

To assess symbiont shifts under future climate scenarios, recruits were subsequently exposed to three experimental treatments representing present-day (hereafter referred to as "ambient") (28.5˚C, $p$CO$_2$ 400 ± 60ppm), 2050 (+1°C offset, $p$CO$_2$ 685±60ppm), and 2100 (+2°C offset, pCO$_2$ 940±60ppm) conditions forecast under 8.5 RCP (Meinshausen *et al.*, 2011, Collins *et al.*, 2013) with replicate tank and treatment conditions as per Botté *et al.* (2020). Control temperature was based on the daily average sea surface temperature from Davies reef in the central GBR (temperature from 1991 to 2012), and control $p$CO$_2$ reflected present-day conditions (Uthicke *et* *al.*, 2020) (Supplementary Table 1). The 6-well plates with coral recruits were randomly distributed across 3 mesocosms per treatment (n=2 plates per mesocosm per species). Well information was not collected for individuals and was therefore not assessed as a factor.

#### C. Sampling early life history stages of corals

Juveniles were sampled directly before adding the 6-well plates to the experimental tanks (hereafter referred to as T0). This was done using a sterile scalpel and juveniles were preserved in absolute ethanol (n=3 individuals per tube). After 10 days (T1) and 4 (*G. retiformis*) or 5 (*A. millepora*) weeks (T2) of exposure to climate conditions, *G. retiformis* and *A. millepora* juveniles were sampled and preserved in absolute ethanol. All samples were stored in -80˚C freezers until further processing.

#### Extractions and sequencing

Genomic DNA was extracted from *G. retiformis* and *A. millepora* samples across three sampling times (T0, T1 and T2) and treatments (ambient, 2050, 2100). Note that the T0 sampling timepoint occurred immediately before juveniles were placed into the three experimental treatments. DNA extractions were performed on individual coral juveniles using a KOH-EDTA method following Sun *et al.* (2014), with increased incubation time (15 minutes at 70˚ C) to enhance cellular lysis.

Polymerase Chain Reaction (PCR) amplification of the ITS-2 locus were conducted using MyTaq DNA Polymerase (Bioline) following manufacturer's protocols with an addition of 1µl 50mM magnesium. The ITS-2 locus was amplified using ITS2alg-F (5'-TCGTCGGCAGCGTCAGATGTGTATAAGAG ACAGGTGAATTGCAGAACTCCGTG) and ITS2alg-R(3'-TTCGTATATTCATTCGCCTCCGACAGAGAATATGTGTAGAGGCTCGGGTGCTCTG-5') primers. PCR was performed on per sample in 25µl reactions (2µL template). PCR conditions were the following: initial denaturation at 95°C for 10 minutes, 32 cycles at 95°C for 30 sec, 59°C for 60 sec, 72°C for 30 sec, and a final elongation at 72°C for 7 minutes. PCR amplification products (see Supplementary Table 2 and 3 for replicate numbers) were delivered to the

Ramaciotti Centre for Genomics (UNSW, Sydney) for Miseq amplicon sequencing of the 300 bp ITS-2 region (LaJeunesse, 2001, LaJeunesse, 2002).

#### D. Bioinformatics and data analysis

A total of 137 individual juvenile samples were successfully sequenced (i.e. some samples were removed if PCRs did not result in bands). Of these successfully sequenced juveniles, 27 samples belonged to *A. millepora*, and 110 belonged to *G. retiformis* (see Supplementary Table 2).

Symbiodiniaceae taxonomy is currently undergoing an extensive revision (LaJeunesse *et al.*, 2018) with substantial effort to understand how sequence diversity links with species, genus, and family designations and most importantly, the functional significance of the symbiont within the coral. Presently, there are generally two analysis pipelines. The first groups sequences within genera (Symportal, Hume *et al*., 2019), and is generally good for high-abundance taxa and making broad inferences across treatments. The other is based on characterizing individual sequence variants using DADA2 (ASVs; Quigley *et al.* 2019; Callahan *et al*., 2016), and provides a higher resolution at the sequence level for exploration of potential diversity at both high and low abundances. The choice of method depends on the question being asked, where both will not render species- or functional- level designations with a formal taxonomic description. Here we chose to use the DADA2 method given our questions about the earliest uptake of symbiont cells and acknowledge that some of the ASVs discovered may be spurious sequence variants and may not represent actual Symbiodiniaceae "species" per-se.

The full pipeline and scripts are described in Quigley et al. 2019 and links therein. Briefly, fastq files were first filtered to remove any reads with retained sequencing adapters using BBDuk (BBMap, package 38.63 http://sourceforge.net/projects/bbmap/) (Bushnell *et al.*, 2017) in R (R Core Team, 2021). Based on quality filtering using BBDuk, reads were then assessed for low quality were trimmed by removing reads without an overlap of 30 bp and one expected error. Dereplication of reads grouped unique sequences, which eliminates any redundant comparisons in the pipeline. Culling spurious sequence variants by merging denoised forward and reverse reads and removing none-overlapping paired reads further helps to remove spurious sequences. Error rates for forward and reverse reads are learnt, both ends are merged and used as input into the DADA2 naive RDP's Bayesian classifier to assign amplicon sequence variants (hereafter ASVs).

After chimera removal, the assigned taxonomy function within DADA2 was used to classify ASVs to known Symbiodiniaceae sequences from the GeoSymbio ITS2 database (Franklin *et al.*, 2012). In essence, this procedure groupings based on sequence similarity, with names derived from the literature as represented in the GeoSymbio database, but do not correspond with strict taxonomic designations, i.e. no binomial species name. Bootstrapping at a threshold of 50 allowed for maximum retention of sequences while minimizing low quality matches. Here we refer to ASVs when discussing the sequence level, and "types" when discussing multiple ASVs that are assigned to the same Symbiodiniaceae taxa (i.e "ASVs" ASV1_C1 and ASV2_C1 are collapsed into "type" C1). On average, 100,801.90 ± 4,212.15 sequences were retained in each sample after quality filtering (see Supplementary Table 2 for summary).

Percent relative abundances of each ASV given the total number of cleaned reads were calculated per sample using the *abundance* (compositional) function in the package "microbiome" (v. 1.10.0)(Lahti *et al.*, 2017). Alpha and beta diversity metrics were calculated using the relative abundance data in "Phyloseq" (v. 1.30.0)(McMurdie & Holmes, 2013).

Alpha diversity was measured using the Shannon index, and linear mixed models were performed to test differences in diversity across time, treatments and species using the R package lmerTest (Kuznetsova *et al.*, 2017). The assumptions of normality, homogeneity of variance and linearity were tested using the Shapiro-Wilk test, Fligner-Killeen test and visual inspection of the residuals (Hartig, 2021). To test for differences in diversity across time (T0, T1, T2) and treatments (Ambient, 2050, 2100) in *G. retiformis*, a linear mixed model that included time and treatment as fixed effects (and their interactions) and tank as random effect was performed. For *A. millepora*, the effect of climate treatment on diversity was tested using a linear mixed model using treatment as fixed factor and tank as random factor on square-root transformed data. To test for differences in alpha diversity between species at T2, a linear mixed model with species and treatment (and their interaction) as fixed factors and tank as random factor was run. Post-hoc multiple comparisons adjusted by the False Discovery Rate (Benjamini & Hochberg, 1995) were run in case of significant interactions (or factors) using the R package multcomp (Hothorn *et al.*, 2008).

To test for significant differences in the relative abundances of each ASV across treatments, the packages DESeq (v. 1.39.0) and DESeq2 (v. 1.26.0) (Love *et al.*, 2014) were used. Generalized linear models with time (T0, T1, T2) and treatment (Ambient, 2050, 2100) as factors were run for 30 iterations, where then non-converging models were filtered prior to multiple test correcting. Significantly differential abundant ASVs were then aligned using Clustal Omega (Sievers *et al.*, 2011).

#### G. Risk Assessment

Symbiodiniaceae 'phylotypes' of interest based on the results of the total abundance, relative abundance, and significant shifts were identified in addition to close relatives based on phylogenic studies. The IPRAPA-based Rk values determined by Swain et al. (2017) measuring thermosensitivity based on experimental studies of shifts in Symbiodiniaceae genera (referred to as 'phylotypes') were used for identification of observed response, labeled as 'Rk (Shift) Score'.

Source of stress was calculated for each type based on 12 measurements (e.g. Fv/Fm, chlorophyll content, lipid assessment) collected from 17 publications (Supplementary Table 3). This calculated the average change across each measurement (e.g. Fv/Fm) for each type (e.g. D1a) and ranked based on the smallest change (-0.1, ranked 1; -0,51 ranked 13), as this would indicate greater resilience to stress and therefore this type being the best performing. Ranks were scored using a Borda Rank method (Borda, J.C. 1781, Roy et al. 2021) and relative rankings were averaged for a final rank score of 1-21 with the smallest score being the 'most important' based on their functional contributions (Supplementary Table 4).

### Supplementary Tables and Figures

**Table 1**: Summary of target temperature / $p\mathrm{CO_2}$ levels at each sampling time (T1, T2) across climate treatments (Ambient, 2050, 2100)

| | Ambient | 2050 | 2100 |
|---|---|---|---|
| *G. retiformis* T1 | 27.9 ˚C / 400 ppm | 28.9 ˚C / 685 ppm | 29.9 ˚C / 940 ppm |
| *G. retiformis* T2 | 28.3 ˚C / 400 ppm | 29.3 ˚C / 685 ppm | 30.3 ˚C / 940 ppm |
| *A. millepora* T2 | 28.6 ˚C / 400 ppm | 29.6 ˚C / 685 ppm | 30.6 ˚C / 940 ppm |

**Table 2**: Summary of samples with successful PCRs and sequencing, processed through the Symbiodiniaceae DADA2 pipeline for ITS2 described in Quigley et al. 2019. Following the cleaning steps, 14 samples were removed

due to low quality or quantity of reads. This included 1 T0 ambient, 3 T0 2050,  5 T1 ambient, 4 T1 2050, and 1 T1 2100, for a remaining sample size of 135.

| ***Tank rep*** | ***Tank Number*** | ***Treatment*** | ***G. ret T0*** | ***G. ret T1*** | ***G.ret T2*** | ***A. mill T2*** |
|---|---|---|---|---|---|---|
| **Replicate 1** | **3** | **Ambient** | **4** | **11** | **11** | **3** |
| **Replicate 2** | **6** | **Ambient** | **3** | **4** | **6** | **3** |
| **Replicate 3** | **8** | **Ambient** | **1** | **3** | **3** | **3** |
| **Replicate 1** | **1** | **2050** | **5** | **4** | **3** | **3** |
| **Replicate 2** | **4** | **2050** | **2** | **3** | **2** | **3** |
| **Replicate 3** | **7** | **2050** | **2** | **6** | **3** | **4** |
| **Replicate 1** | **2** | **2100** | **4** | **7** | **1** | **2** |
| **Replicate 2** | **5** | **2100** | **3** | **6** | **0** | **3** |
| **Replicate 3** | **9** | **2100** | **1** | **5** | **7** | **3** |

**Table 3**: Redundancy scores were calculated using results of experimental studies compiled from 17 publications which measured for a variety of 12 measurements in corals and their endosymbionts. Redundancy scores were ranked using a Borda-rank method for those which presented the smallest change. A smaller shift was assumed to be indicative of greater resilience to stress and therefore higher performing in that function.

| Source | Measurements | Species | Symbiont |
|---|---|---|---|
| **de Luna 2020** | Fv/Fm (PSII)<br>chlorophyll content (Chl a + *c2*)<br>PSI:PSII<br>Dark oxygen consumption (Rd) | S. pistillata | Durusdinium |
| **Gong 2019** | Fv/Fm (PSII) | P. lamellina<br>P. lutea<br>F. speciosa | C21<br>C3b<br>C3<br>D1<br>C15<br>C91<br>C3u |
| **Hoadley 2019** | Fv/Fm (PSII)<br>Chlorophyll Content (Chl a + *c2*) | P. rugosa<br>A. muricata<br>C. chalcidicum<br>Coelastrea<br>P. rugosa<br>A. muricata<br>C. chalcidicum<br>Coelastrea | D1a<br>C40<br>C41<br>C21<br>C22<br>C3u |

| | | | |
|---|---|---|---|
| **Gardner 2017** | Fv/Fm (PSII)<br>Chlorophyll Content (Chl a + *c2*)<br>Photosynthesis:Respiration<br>DMSP concentration<br>DMSO<br>Glutanthione<br>Catalase-like activity<br>Superoxide dismutase<br>Light Pressure | A. millepora<br>S. pistillata | C3<br>C8a |
| **Krueger 2015** | Fv/Fm (PSII)<br>chlorophyll content (Chl a + *c2*)<br>Glutanthione<br>Catalase-like activity<br>Superoxide dismutase | A. millepora<br>M. digitata | C3<br>C15 |
| **Shore-Maggion 2018** | Fv/Fm (PSII)<br>Bleaching response | red M. capitata<br>orange M. capitata | C3<br>D1<br>C1 |
| **Fischer 2011** | Fv/Fm (PSII)<br>Light Pressure | A. millepora<br>P.daedalea<br>A. aspera<br>A. formosa<br>M. digitata<br>P. cylindrica<br>P. lutea<br>M. digitata | C3<br>C15<br>C17* |
| **Kenkel 2018** | Rate of photosynthesis | A. millepora<br>M. aequituberculata<br>P. lobata<br>G. columna<br>G. astrea<br>G. acrhelia | C15<br>D1<br>C3<br>C35<br>C15<br>C1<br>Cspc<br>D1a<br>D6<br>C35 |
| **Abrego 2008** | Fv/Fm (PSII)<br>Chlorophyll Content (Chl a + *c2*)<br>Rate of Photosynthesis<br>Bleaching response | A. tenuis | C1<br>D1a |
| **Schoepf 2015** | Experiment type<br>Fv/Fm (PSII)<br>Qm - Maximum excitation pressure<br>Chlorophyll Content (Chl a + *c2*) | A. aspera<br>Dipastrea sp. | Cp1 |

| | | | |
|---|---|---|---|
| **Ros 2021** | Dark oxygen consumption (Rd)<br>Photosynthesis:Respiration<br>Rate of Photosynthesis | P. acutata | C1<br>D6<br>C1b<br>D1-D4-D2.2 |
| **Swain 2016** | Fv/Fm (PSII)<br>Qm - Maximum excitation pressure | Merlina sp.<br>P. damicornis<br>S. hystrix<br>S. pistillata<br>T. reniformis<br>Goniapora sp.<br>F. favus<br>M. foliosa<br>M. digitata | C3u<br>D1<br>D1a<br>C8a<br>C3v<br>C3u<br>C15 |
| **Silverstein 2012** | Cladal function - thermosensitivity<br>Cladal function - growth<br>Cladal function - Photochemical efficiency<br>Fv/Fm (PSII) | | Cladocopium |
| **Silverstein 2011** | Cladal function - thermosensitivity<br>Cladal function - growth<br>Cladal function - Photochemical efficiency<br>Fv/Fm (PSII) | | Durusdinium<br>Cladocopium |
| **Aschaffenburg 2012 Diss.** | Cladal function - thermosensitivity<br>Cladal function - growth<br>Cladal function - Photochemical efficiency | | Cladocopium<br>Durusdinium |
| **Kneeland 2013** | Cladal function - thermosensitivity<br>Cladal function - growth<br>Cladal function - Photochemical efficiency<br>Fv/Fm (PSII) | | Durusdiniu |
| **Morgans 2019** | Cladal function - thermosensitivity<br>Cladal function - growth<br>Cladal function - Photochemical efficiency<br>Fv/Fm (PSII) | | Durusdinium |

**Table 4**: Final risk assessment plotting is based on Rk values (*sensu* Swain et al. 2017) as well as final Redundancy rankings from a Borda rank-based assessment of functional redundancy of the 16 Symbiodiniaceae species of interest.

| Phylotype | Redundancy Rank | Rk (Shift) Value | Relative |
|---|---|---|---|
| C1 | 8 | 21.72 | |
| C1b | 6 | 19.11 | |
| C1c | 6 | 35.71 | |
| C1m | 0 | 5.27 | |
| C3 | 11 | 25.98 | C16a |

| | | | |
|---|---|---|---|
| C3u | 16 | 21.95 | |
| C3v | 12 | 0 | |
| C15 | 15 | 53.12 | |
| C17* | 10 | 10.62 | |
| C21 | 3 | 40.25 | |
| C35 | 17 | 0 | |
| C40 | 7 | 55.32 | |
| D1 | 5 | 43.82 | |
| D1a | 13 | 37.03 | D6 |
| D4 | 4 | 4.39 | D9 |
| D6 | 1 | 0 | |
